# Intraspecific genetic variation underlying postmating reproductive barriers between species in the wild tomato clade (*Solanum* sect. *Lycopersicon*)

**DOI:** 10.1101/718544

**Authors:** Cathleen P Jewell, Simo Zhang, Matthew J. S. Gibson, Alejandro Tovar-Méndez, Bruce McClure, Leonie C Moyle

## Abstract

A goal of speciation genetics is to understand how the genetic components underlying interspecific reproductive barriers originate within species. Unilateral incompatibility (UI) is a postmating prezygotic barrier in which pollen rejection in the female reproductive tract (style) occurs in only one direction of an interspecific cross. Natural variation in the strength of UI has been observed among populations within species in the wild tomato clade. In some cases, molecular loci underlying self-incompatibility (SI) are associated with this variation in UI, but the mechanistic connection between these intra- and inter-specific pollen rejection behaviors is poorly understood in most instances. We generated an F_2_ population between SI and SC genotypes of a single species, *Solanum pennellii*, to examine the genetic basis of intraspecific variation in the strength of UI against other species, and to determine whether loci underlying SI are genetically associated with this variation. We found that F_2_ individuals vary in the rate at which UI rejection occurs. One large effect QTL detected for this trait co-localized with the SI-determining *S*-locus. Moreover, individuals that expressed S-RNase—the *S-*locus protein involved in SI pollen rejection—in their styles had much more rapid UI responses compared to those without S-RNase protein. Our analysis shows that intraspecific variation at mate choice loci—in this case at loci that prevent self-fertilization—can contribute to variation in the strength of interspecific isolation, including postmating prezygotic barriers. Understanding the nature of such standing variation can provide insight into the accumulation of these barriers between diverging lineages.

## Introduction

Speciation involves the accumulation of genetic differences and—in sexually reproducing organisms—reproductive isolation, among diverging lineages. Accordingly, loci that contribute to this cumulative process between species must first arise within an individual population prior to spreading to other conspecific populations within their own lineage. During this process, populations of a single species are expected to show varying strengths of reproductive isolation against other lineages; that is, there will be intraspecific genetic variation for the magnitude of interspecific reproductive isolation from other lineages. Intraspecific phenotypic variation in the strength of hybrid incompatibility has been observed in many systems including mammals (Good, Handel and Nachman, 2007; Vyskočilová, Pražanová and Piálek, 2009), arthropods (Bordenstein, Drapeau and Werren, 2000; Kopp and Frank, 2005; Reed and Markow, 2004; Shuker, *et al*., 2005), nematodes (Kozlowska, *et al*., 2012) and plants (Case and Willis, 2008; Leppälä and Savolainen, 2011; Martin and Willis, 2010; Rieseberg, 2000; Sweigart, Mason and Willis, 2007). Understanding the nature, origin, and accumulation of this variation, including the underlying molecular genetic variants responsible, can provide insight into the evolutionary dynamics of lineage divergence (Cutter, 2012), including the order in which alleles contributing to interspecific reproductive isolation arise and fix within diverging lineages.

The genetic basis of intraspecific variation for interspecific barriers has been investigated in few cases, most of which focus on postzygotic isolating barriers. Some of these studies have confirmed that variable reproductive isolation is due to genetic variation between populations of a species, but have not identified the specific loci or gene(s) responsible [e.g. (Kozlowska, *et al*., 2012; Machado, Haselkorn and Noor, 2007)]. In other cases, isolation variation has been mapped to localized chromosomal regions (quantitative trait loci, or QTL) or even individual loci, whose geographic distribution is then investigated. For example, between the plant sister species *Mimulus nasutus and M. guttatus*, the *M. guttatus hybrid male sterility 1* (*hms1*) allele interacts with the *M. nasutus hms2* allele to cause male sterility in hybrids; the *M. nasutus hms2* allele is common across populations, but the interacting *hms1* allele is geographically restricted within *M. guttatus* (Martin and Willis, 2010; Sweigart, *et al*., 2007; Zwellig and Sweigart 2018a, b). This and other studies of hybrid sterility and inviability (e.g., Leppälä and Savolainen, 2011; Reed and Markow, 2004; Shuker, *et al*., 2005) confirm that variation in the strength of interspecific isolation between populations within species can be due to standing genetic variation.

Compared to these studies of variable postzygotic isolating barriers, fewer analyses address within-species polymorphism for prezygotic reproductive isolation [for an exception, see (Hopkins and Rausher, 2012)]. Because prezygotic barriers act earlier in reproduction, they could have a much larger role than postzygotic barriers in restricting levels of gene flow between closely related species (Lowry, *et al*., 2008; Rieseberg and Willis, 2007). Of these, postmating prezygotic interactions can be particularly important for reproductive isolation when species are only weakly isolated by other prezygotic mechanisms, such as pollinator isolation [reviewed in (Swanson, Edland, and Pruess 2004; Moyle, Jewell and Kostyun, 2014)].

Unilateral incompatibility (UI) is an example of a postmating prezygotic isolating barrier that shows variation in strength among populations within species. In plants, this barrier manifests after pollen transfer, as the (male) pollen grains germinate and produce pollen tubes that grow down the female reproductive tract (the ‘pistil’, composed of the stigma (the pollen receiving site), the ovary, and the style which connects them) towards individual ovules. UI occurs between species when pollen rejection in the female style occurs in only one direction of an interspecific cross (and is therefore ‘unilateral’; de Nettancourt, 1977; Lewis and Crowe, 1958), while the reciprocal cross results in pollen tubes successfully growing down the style and into the ovary. UI often follows an ‘SI x SC rule’ in which genetically self-incompatible (SI) species reject pollen from self-compatible (SC) species but the reciprocal cross is successful (de Nettancourt, 1977; Lewis and Crowe, 1958; Murfett, *et al*., 1996); however, there are exceptions to this rule even within species (Baek, *et al*., 2015). In the wild tomato clade, species that are largely SI and display strong UI responses can nevertheless include SC populations that exhibit weakened UI. In some species, the UI response is less rapid in SC populations, but in other species, SC populations fail to reject heterospecific pollen altogether (i.e. UI is lost in these populations) (Baek, *et al*., 2015, and see Discussion).

These observations suggest that among-population variation in the strength of interspecific UI might be mechanistically associated with molecular factors contributing to SI (Li and Chetelat, 2015; Tovar-Méndez, *et al*., 2014). However, the extent to which UI and SI are consistently genetically associated remains unclear. Within the plant family Solanaceae, a primary determinant of gametophytic self-incompatibility is the *S-*locus (McClure, Cruz-Garcia and Romero, 2011; McClure, *et al*., 1989) which encodes at least two proteins responsible for the self-rejection mechanism: an S-RNase protein (the stylar component) that recognizes one or more pollen-expressed F-box protein(s) in germinated pollen tubes and arrests pollen tube growth within the style (Kubo, *et al*., 2015; Kubo, *et al*., 2010; Li and Chetelat, 2015; Sijacic, *et al*., 2004; Williams, *et al*., 2014). Pollen is rejected when a haploid pollen tube bears an S-haplotype that is identical one of the *S*-haplotypes of the pistil (maternal) parent (McClure, *et al*., 1989). Loss of SI in the wild tomato clade is frequently associated with mutations in the locus producing *S-RNase* (Bedinger, *et al*., 2011; Igic, Lande and Kohn, 2008; Rick and Chetelat, 1991). Similarly, within predominantly SI species, population-level transitions to SC are also often associated with the loss of *S-RNase* function. Within the two wild tomato species *S. habrochaites* and *S. pennellii*, for example, several SC populations have been shown to lack *S-RNase* expression in stylar tissue (Kondo et al. 2002). Despite the loss of *S-RNase*, however, most SC populations of these wild species still exhibit UI against interspecific pollen, indicating UI can also have S-RNase-independent mechanisms. Nonetheless, the degree to which natural intraspecific variation in genes involved in SI might simultaneously affect interspecific isolation via UI remains to be directly investigated in most cases (but see Broz et al. 2017, Markova et al. 2016).

In this study, we generated an F_2_ mapping population between two populations within a single species to map QTL underlying variation in the strength of UI against a second, tester, species. Our two parental genotypes were drawn from an SI population (*S. pennellii* accession LA3778) and a conspecific SC population (*S. pennellii* accession LA0716) which has recently lost SI; both these genotypes express UI, but differ in how fast they reject pollen from domesticated tomato pollen and other SC species (*Results*). Our goal was to determine the genetic basis of this standing variation for the rate of UI rejection within our target species, and its association with molecular loci underlying SI. We quantified UI response, evaluated SI status, and measured several floral and fertility traits in the recombinant F_2_ population. We assessed 1) the number of large effect QTL that contribute to variation in the UI response within *S. pennellii*, 2) the association, if any, between these UI loci and *a priori* candidate loci known to contribute to intraspecific SI variation, and 3) the degree of association, if any, between UI phenotypes and floral or fertility traits. These data allow us to assess whether the strength of UI (and changes in SI status) in different intraspecific *Solanum* lineages is due to changes at the same underlying loci.

## Materials and Methods

### Generating the F_2_ population

The wild tomato clade, *Solanum* sect. *Lycopersicon* is a group within the diverse nightshade family *Solanaceae* that consists of 13 closely related [<2.5 MY old; (Pease, *et al*., 2016; Peralta, Spooner and Knapp, 2008; Rodriguez, *et al*., 2009)] hermaphroditic species, including the domesticated tomato and its wild relatives (Peralta, *et al*., 2008). In this study our focal species was *Solanum pennellii*, a wild, herbaceous, perennial species (**Supplementary Figure 1**). *S. pennellii* populations--including the two parental accessions (populations) used here (see below)--can vary in the strength of UI against other SC *Solanum* species (Liedl, McCormick and Mutschler, 1996). We generated a recombinant F_2_ population in which the female parent was from self-compatible *S. pennellii* accession LA0716, and the male parent was from self-incompatible *S. pennellii* accession LA3778. LA0716 does not express *S-RNase*, likely due to a deletion in the underlying gene (Li and Chetelat, 2015), however both accessions exhibit UI against other SC species, including domesticated tomato.

Seeds of the parental accessions were obtained from the Tomato Genomics Resource Center (TGRC; tgrc.ucdavis.edu), grown to maturity and one individual from each accession was chosen to make the original cross. The F_1_ offspring of this cross were self-compatible and one F_1_ was selfed by hand-pollination to generate the F_2_ generation (n = 100). To cultivate all experimental plants, seeds were treated with 50% bleach for 30 minutes, rinsed, placed on moist blotting paper and incubated (12hr day-length, 24°C) to stimulate germination. Germinated seedlings were transplanted into flats with Metro Mix 360 (Sun Gro) potting mix and hand watered daily. Once well-established, the seedlings were transferred to individual 1-gallon pots containing 50% Metro Mix 360 and 50% Indiana University (IU) greenhouse potting mix; pots were placed in a climate controlled greenhouse at IU with 14hr day-length. Plants were watered twice daily, fertilized weekly, staked prior to flowering, and regularly pruned thereafter.

### Quantifying unilateral incompatibility

To assess the quantitative expression of interspecific UI, each F_2_ was pollinated with the same tester genotype of *S. lycopersicum* (accession LA3475, SC). While both SC LA0716 and SI LA3778 reject *S. lycopersicum* pollen, the former has a slower UI response (i.e. the pollen is halted after growing further down the style; *Results*). At least 3 flowers from each F_2_ individual were emasculated one day prior to anthesis, the styles pollinated 24hrs later, and collected after an additional 24hrs, which is sufficient time for compatible (i.e., conspecific) pollen tubes to reach the ovary in the parental genotypes. Styles were fixed in 3:1 ethanol:glacial acetic acid, stained using aniline blue fluorochrome (Biosupplies Australia Pty Ltd) and imaged using fluorescent microscopy. Because styles were too long to be captured in one image frame, several images were taken along the axis of each style and then stitched (Autostitch; (Brown and Lowe, 2007)). Stitched images were visualized for data collection using ImageJ (Abramoff, Magalhaes and Ram, 2004). UI response (location of pollen rejection within the style) was quantified by measuring the total style length, length of the five longest pollen tubes, and length of the pollen tube “front” where the majority of pollen tubes stopped growing. In all analyses reported here, UI response was calculated by dividing the average of the five longest pollen tubes by the total length of the style. Thus, mean pollen tube growth is quantified as a proportion of style length travelled and varies from 0 (representing no growth down the style) to 1 (where pollen tubes reach the end of the style). UI was similarly quantified in the parents and F_1_ as described above.

### Evaluating self-incompatibility status

In gametophytic SI, complete pollen rejection only occurs if both *S*-haplotypes in the pistil match both *S*-haplotypes in the pollen parent. Because our F_2_ population was generated by self-fertilization of one F_1_ individual, a simple SI/SC nomenclature cannot be applied to this population. Instead, we expected that F_2_s would display ‘acceptor’ phenotypes, as they would have at most one functional *S-*haplotype from the original SI parent. For example, here, if the LA3778 parent is designated *S*_1_*S*_2_ and the LA0716 parent *S*_0_*S*_0_, their F_1_ could be *S*_1_*S*_0_ or *S*_2_*S*_0_. During self-fertilization of a particular F_1_ individual (e.g., *S*_1_*S*_0_), pollen bearing the LA3778 (SI parent) *S*_1_-haplotype will be selectively arrested, leaving only pollen with LA0716 (SC parent) *S*_0_-haplotype to fertilize the F_1_ ovules; the resulting F_2_ individuals are therefore expected to be either *S*_1_*S*_0_ or *S*_0_*S*_0_, and none should completely reject pollen from either SI or SC parent (i.e. they are ‘acceptors’). To confirm this was the case, we evaluated the pollen-rejection status of individuals in several ways. To initially test self-fertility status, at least three flowers from each F_2_ individual were manually self-pollinated. Selfed F_2_s that produced fruits were designated as acceptor phenotypes. Fruits from these pollinations were left to mature on the plant; at maturity each was weighed and measured (length and width), and seeds extracted by hand to count viable seeds per fruit. In the rare cases (i.e., three individuals) where no fruits were produced from these initial hand-pollinations, individuals were provisionally designated as self-sterile. These individuals were further evaluated using pollen from the original SI LA3778 parent used to generate the F2, first by evaluating fruit set following pollination and then by directly assessing pollen rejection by visualizing pollen tube growth in styles. The latter experiments were performed as for UI (see ‘Quantifying unilateral incompatibility’ above), except that the tester pollen came from the LA3778 parent. The F_1_ was evaluated for self-fertility/acceptor status in the same manner.

### Floral morphology and fertility traits

To quantify additional reproductive traits that might be relevant to mating system variation or to the expression of UI, six floral and six fertility traits were measured. The six floral traits were: corolla diameter, style length, stigma exsertion (distance between stigmatic surface to the tip of the anther cone, on an intact flower), anther length, ovary height, and ovary width [(Moyle, 2007); **Supplementary Figure 1**]. Using digital calipers, three fully open flowers (day 1 of opening) per F_2_ individual were measured for all floral traits, and replicate measures averaged within each individual prior to analyses. For each parent individual and the F_1_, five replicate flowers were similarly measured. For overall comparison of the two parental accessions, floral traits were also quantified on five additional individuals from each accession, by taking the average of measurements from three flowers per individual.

For fertility traits, we quantified total pollen, proportion viable pollen, fruit weight, fruit width, fruit length, and seed set. Pollen number per flower was estimated by collecting whole anther cones from individual flowers one day before opening, into lactophenol aniline blue histochemical stain (Kearns and Inouye, 1993; Moyle and Graham, 2005). Each anther cone was homogenized and for each sample an aliquot of homogenate was examined on an inverted microscope using a hemacytometer, to count total pollen grains and estimate proportion of viable and inviable pollen. Pollen that fails to stain lacks functional cytoplasm and was classified as inviable. At least three anther cones were collected and counted per individual; mean counts for each individual were used for analysis. At least three selfed fruits per individual (where possible) were hand harvested, individually weighed, and bisected to take length and width measurements (see ‘Self-incompatibility status’ section above). All seeds were extracted and the number of viable seeds was counted per fruit. At least three fruit were measured per individual and trait means for each individual were used for analysis.

### Style Protein Expression

Two loci (*S-RNase* and *HT*) were directly investigated for their association with UI phenotypic variation, by assessing their protein expression in parental, F1, and F2 styles. Both S-RNase protein and HT (which is a small asparagine-rich protein (Covey, *et al*., 2010; McClure, *et al*., 1999; O’Brien M, 2002)) have been previously implicated in UI expression (Murfett *et al*., 1996, Tovar-Mendez *et al*., 2014)—including in QTL mapping studies in other *Solanum* species (Bernacci and Tanksley, 1997)—making them *a priori* candidates for UI variation in this population. Both genes are also essential for SI (McClure, *et al*., 1999). The *HT* gene was duplicated in the ancestor of *Solanum*, giving rise to two tandemly arrayed genes (Sopen12g029190, *HT-A* and Sopen12g029200, *HT-B*) on chromosome 12 (Covey et al. 2010). A subset of individuals (F_1_ n = 1; F_2_ n = 21) was screened for protein expression of S-RNase and HT using protein blotting. For each individual, flowers were emasculated 24hrs before opening; styles were collected 24hrs later and weighed. At least 5mg stylar tissue was collected per individual and protein was extracted using 2xLSB (Laemmli Sample Buffer; 10uL LSB / 1mg tissue). Samples were boiled 5min, centrifuged (10min at 20,000 x g) and the supernatant was retained for analysis. For S-RNase detection, extract equivalent to 0.2mg fresh weight per lane was separated in 10% Tris-Tricine SDS PAGE, blotted to PVDF, and immunostained (1:5000) with an antibody against the conserved C2 S-RNase motif, as described previously (Covey, *et al*., 2010). For HT-protein detection, extracts equivalent to 1.5mg fresh weight were separated in 12.5% Tris-Tricine SDS PAGE, blotted to PVDF, and immunostained (1:5000) with an affinity-purified antibody that recognizes both HT-A and HT-B proteins. The antibody was prepared against the synthetic peptide LEANEIHNTELNNPTLQKKGGC-amide (21st Century Biochemicals), as described previously (Tovar-Méndez, *et al*., 2017).

### Genotyping F_2_s

To genotype our F_2_ population, genomic DNA from 93 F_2_s and each parent was extracted using Qiagen DNeasy Plant Mini Kits. Extracted genomic DNA samples were sent to the Cornell University Institute of Biotechnology’s Genomic Diversity Facility for genotyping-by-sequencing (GBS), using restriction enzyme PstI. An unfiltered SNP marker set was generated by the Cornell Institute of Biotechnology, by mapping trimmed raw sequence reads onto the *S. pennellii* (LA0716) genome (Bolger, *et al*., 2014) using *bwa* (Li 2013), within the Cornell *TASSEL* 3.0 GBS reference pipeline (version 3.0.173). To obtain a high-quality set of markers for the linkage map and QTL mapping, only markers with bi-allelic sites and that had fixed differences between parental genotypes were used. For consistency, we required that a maximum of 30% individuals differ in genotypes for any pair of markers that are within 500bp of each other, for these markers to be retained. Segregation distortion was assessed by testing for Hardy-Weinberg equilibrium at each marker, and markers showing significant deviation (p < 0.05) were removed from the final marker dataset. After implementing these filters and also removing samples with low sequencing quality (those that had more than 15% missing genotypes), the initial genetic map contained 810 markers and was significantly expanded (average LG length of 332 cM) likely due to unaccounted for genotyping errors. To accommodate this, we used the Genotype-Corrector tool (Miao et al., 2018) which corrects or imputes genotype calls at reference-mapped markers based on a sliding-window algorithm, prior to rebuilding the linkage map. The resulting dataset contained 569 high-quality markers from 88 individuals; five additional individuals were removed due to high levels of missing data (>15%) following correction or removal of unlikely genotype calls.

### Linkage Map and QTL Mapping

The linkage map was constructed using the Rqtl (Broman, *et al*., 2003) and ASMap (Taylor & Butler, 2017) packages in R version 3.2.2 (R Core Team, 2015); ASMap implements the minimum spanning tree (MST) algorithm (Wu et al., 2008) for map construction. Markers were first clustered by chromosome (based on reference mapping) prior to inferring the marker order on each group using the MST algorithm. The final map length was 1750.48cM (average of 145.87 cM per LG), with an average of 0.276 cM between markers.

For phenotypes that were non-normally distributed (Shapiro-Wilk test; W < 0.05), we transformed the trait data using the nqrank function which transforms the vector of quantitative values to corresponding normal quantiles and preserves the mean and standard deviation. Missing genotypes were imputed prior to performing genome scans with the multiple QTL model [MQM; (Arends, *et al*., 2010)] for each trait. Genome-wide significance LOD thresholds were calculated for each trait based on permutation tests (1000 iterations) with alpha = 0.05. For each trait, we included the significant QTL in a model as the main effect to obtain estimates of the total phenotypic variation explained and individual contributions of each QTL, as well as to test for interactions between QTL. The percent parental difference explained (relative homozygous effect, RHE) was calculated for each detected QTL as the additive QTL effect size divided by the parental difference. We assessed overlap of identified QTL with previously identified UI QTL (Bernacci and Tanksley, 1997) by using information on physical location of markers, the annotated *S. lycopersicum* and *S. pennellii* genomes, and other gene position data from the Sol Genomics Network (solgenomics.net). Finally, we quantified the number of loci that fell within the 1.5-LOD confidence interval (CI) of our UI QTL (see Results) by identifying the two markers closest to each end of the CI and then counting all annotated genes that fell between these two markers, using version 2 of the AUGUSTUS annotation of the *S. pennellii* genome.

### Statistical Analyses

All analyses were run in R version 3.2.2 (R Core Team, 2015) and statistical significance was reported if p < 0.05. Shapiro-Wilks tests were performed to test for normality for each quantitative trait. T-tests were used to test for trait variation between the parental accessions, and to compare UI responses between individuals found to express S-RNase protein or not.

## Results

### Rapidity of unilateral incompatibility rejection response varies among F_2_s

The two parental accessions used to make our F_2_ population both exhibited UI but differed quantitatively in how rapidly they rejected *S. lycopersicum* pollen, i.e. in the position within the style that their UI response manifested. The SI LA3778 parent had a rapid UI response to heterospecific pollen tubes (0.038 ± 0.005 proportion of style length; **Figure 1**), where pollen rejection was defined in terms of the proportional distance of pollen tube growth down the style (so that values closer to zero indicate a rapid UI response whereas slower responses have values closer to 1). The SC LA0716 parent had a less rapid response (0.32 ± 0.16 proportion of style length). The F_1_ individual also expressed a rapid UI response (0.040 ± 0.034 proportion of style length). All measured (n=99) F_2_ individuals expressed UI. Nonetheless, there was broad quantitative variation in the rapidity of the UI rejection response, with the tester pollen tube growth response ranging from 0.01 – 0.55 of the length of F_2_ styles (**Figure 1**).

**Figure 1.**
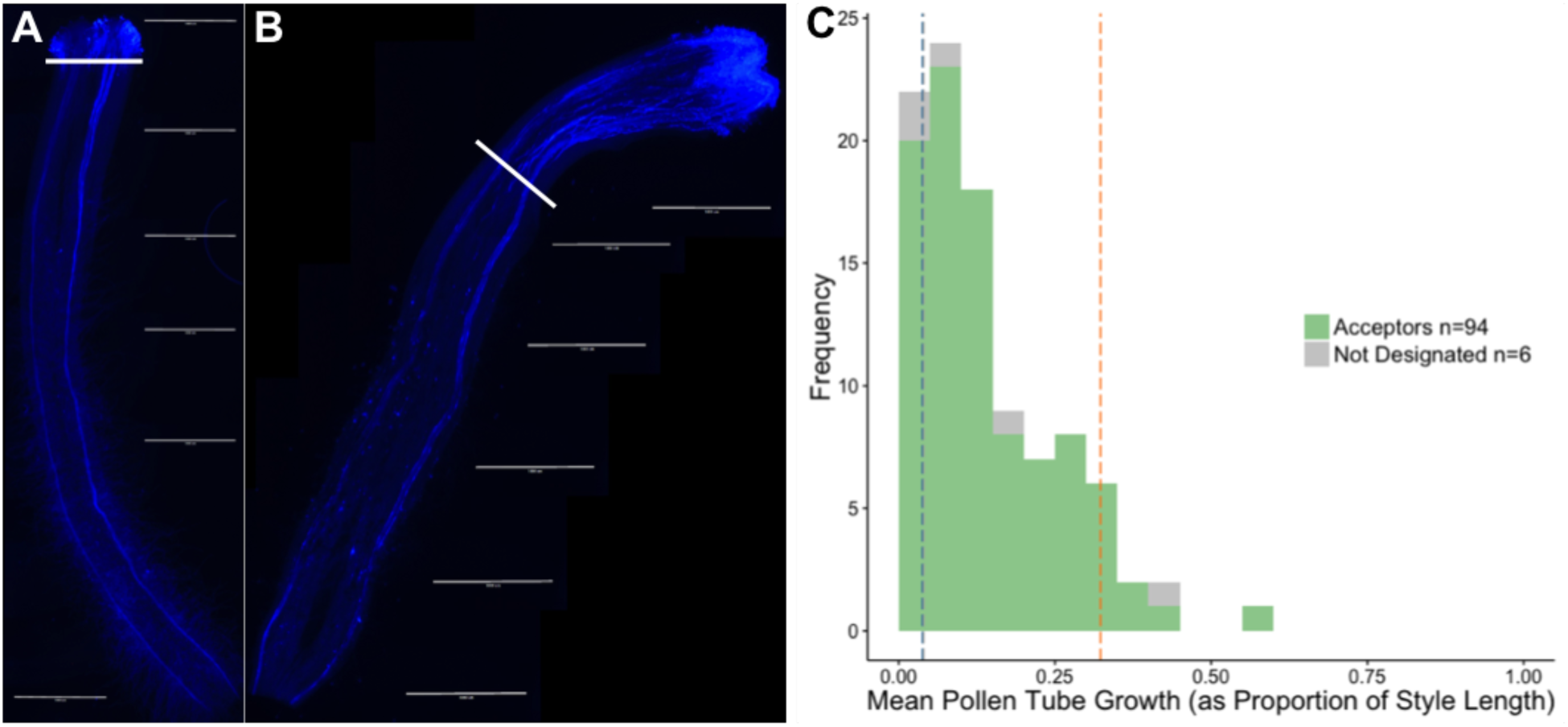
Variation in the strength of unilateral incompatibility (UI) responses between *S. pennellii* populations. **A**. Representative image of an F_2_ with a rapid UI response. **B**. Image of an F_2_ with a less rapid UI response. **C**. Phenotypic distribution of unilateral incompatibility responses across the F_2_ population. Blue dashed line (0.038) is the *S. pennellii* SI parent (LA3778); orange dashed line (0.32) is the *S. pennellii* SC parent (LA0716).

### All F_2_s accepted self-pollen

Because the F_2_ population was generated by self-fertilization of one F_1_ individual, we expected that all F_2_s would accept self-pollen. This is because only F_1_ pollen bearing the LA0716 (SC parent) *S-*haplotype would have fertilized F_1_ ovules (due to pistil-mediated gametophytic selection against the alternative, functional, pollen-side allele from the LA3778 SI parent) (see Methods). We confirmed that all evaluated F_2_ individuals (n = 94) accepted self-pollen (‘acceptors’ in **Figure 1**). In the few individuals (n = 3) that did not successfully develop fruits after self-fertilization, when pollen rejection was directly assessed in their styles, they showed no evidence of a pollen rejection response; that is, pollen tubes were observed growing all the way to the ovary. Therefore we infer that other downstream factors (e.g. low ovule fertility, gamete isolation, or early fruit abortion) may have prevented fruit development in these three individuals.

### Large effect unilateral incompatibility QTL is associated with the S-locus and variation in functional S-RNase

One large effect QTL was identified for UI, explaining 32.6% of the phenotypic variance among F2s, and 23.0% of the difference observed between the parents (**Table 2, Figure 2**). Located on chromosome 1, this QTL colocalizes both with the location of a UI QTL previously identified in a different *Solanum* cross (Bernacci and Tanksley, 1997) and with the genomic location of the *S-*locus (**Figure 2**), which contains genes encoding S-RNase, F-box proteins, and other factors involved in self-incompatibility (Bernacci and Tanksley, 1997; Li and Chetelat, 2015). The 1.5 LOD CI of this QTL spans 29.47cM or ∼85.08Mb, and contains 2684 gene models, based on the v2 annotation of the *S. pennellii* genome. This large low-recombination region is characteristic of the physical location of the *S*-locus (**Figure 2**), which exhibits suppressed recombination in this and other species.

**Table 1.** Trait differences between individuals (n = 5) of the parental accessions, reported as means and standard deviations. T-tests (one-sided) are reported for differences between the parental accessions. Comparisons with p < 0.05 are denoted in bold. F_1_ hybrid values are also reported; NA = not evaluated.

| Trait | *S. pennellii*<br>(LA0716, SC) | *S. pennellii*<br>(LA3778, SI) | t-test | $F_1$ |
| --- | --- | --- | --- | --- |
| UI | $0.322 \pm 0.026$ | $0.038 \pm 2.66 \times 10^{-5}$ | $t = -3.34$; **$p = 0.022$** | $0.039 \pm 3.43 \times 10^{-4}$ |
| Corolla Diameter (mm) | $21.22 \pm 1.88$ | $20.73 \pm 0.86$ | $t = -0.54$, $p = 0.612$ | $28.31 \pm 1.31$ |
| Anther Length (mm) | $7.08 \pm 0.85$ | $7.59 \pm 0.33$ | $t = 1.26$, $p = 0.263$ | $9.44 \pm 0.55$ |
| Stigma Exsertion (mm) | $1.50 \pm 0.28$ | $2.32 \pm 0.43$ | $t = 3.57$, **$p = 0.0096$** | $2.32 \pm 0.52$ |
| Style Length (mm) | $8.29 \pm 1.17$ | $10.56 \pm 0.87$ | $t = 3.48$, **$p = 0.0095$** | $12.23 \pm 0.36$ |
| Ovary Height (mm) | $1.63 \pm 0.12$ | $1.11 \pm 0.05$ | $t = -8.87$, **$p = 0.0003$** | $1.32 \pm 0.13$ |
| Ovary Width (mm) | $1.05 \pm 0.04$ | $0.92 \pm 0.05$ | $t = -4.68$, **$p = 0.002$** | $1.32 \pm 0.05$ |
| Total Pollen | $69.53 \pm 192.8$ | $62.40 \pm 731.3$ | $t = -0.52$; $p = 0.3070$ | NA |
| Proportion Viable Pollen | $0.469 \pm 0.017$ | $0.834 \pm 0.003$ | $t = 5.70$; **$p = 0.0002$** | NA |

**Table 2.** QTL associated with unilateral incompatibility, floral traits, and fertility traits. Percent phenotypic variance (PVE) and percent parental difference (RHE) explained are reported for each trait, with full models including all QTL found for each phenotype.

| Trait | LOD Threshold<br>alpha = 0.05 | QTL peak chromosome, position (cM), closest marker | Peak LOD | 1.5-LOD CI | PVE (%) | Additive effect (S.E.) | Dominance effect (S.E.) | % parental difference explained (RHE) |
| --- | --- | --- | --- | --- | --- | --- | --- | --- |
| Unilateral Incompatibility | 2.71 | 1, 60<br>chr1_79110602 | 5.49 | 40-64.67 | 32.63 | -0.066<br>(0.020) | -0.062<br>(0.024) | 23.0 |
| Corolla Diameter | 2.19 | None |  |  |  |  |  |  |
| Anther Length | 2.85 | None |  |  |  |  |  |  |
| Stigma Exsertion | 2.32 | 4, 180,<br>chr4_75727601 | 2.65 | 75-181.6 | 17 | 0.366<br>(0.082) | 0.092<br>(0.110) | 48.0 |
| Style Length | 2.45 | None |  |  |  |  |  |  |
| Ovary Width | 2.44 | None |  |  |  |  |  |  |
| Ovary Height | 2.19 | None |  |  |  |  |  |  |
| Total Pollen | 2.25 | None |  |  |  |  |  |  |
| Proportion Viable Pollen | 2.12 | None |  |  |  |  |  |  |
| Fruit Width | 2.80 | None |  |  |  |  |  |  |
| Fruit Height | 2.46 | 8, 10,<br>chr8_2708030 | 2.69 | 0-25 | 16.18 | -0.537<br>(0.149) | 0.832<br>(0.271) | 5.0 |
|  |  | 9, 90,<br>chr9_5441571 | 2.68 | 62.19-100 | 6.17 | 0.535<br>(0.173) | -0.047<br>(0.248) | -5.0 |
| Fruit Weight | 2.87 | None |  |  |  |  |  |  |
| Seed Count | 2.55 | None |  |  |  |  |  |  |

**Figure 2.**
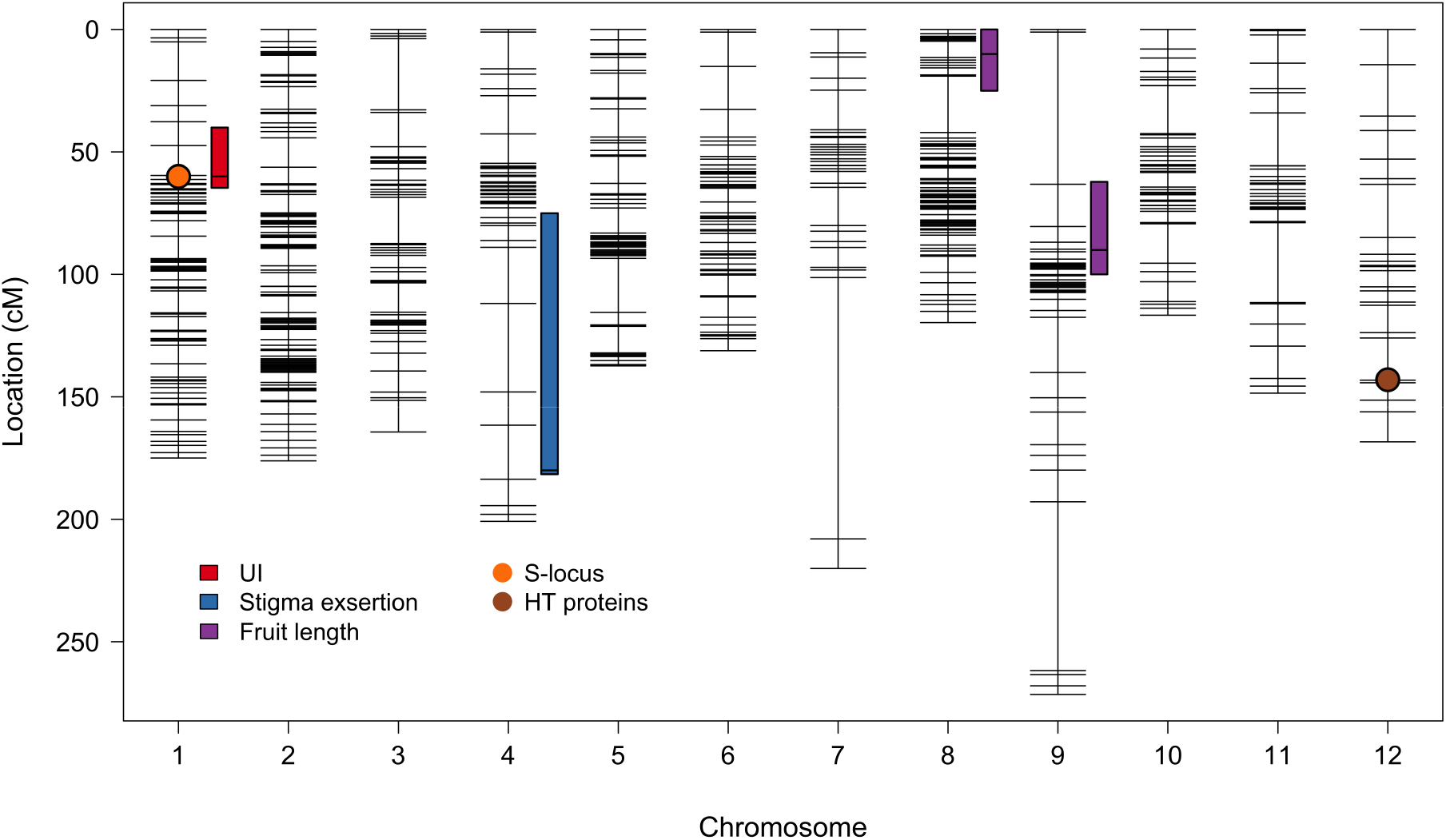
Map of identified QTL. Each QTL is marked with the 1.5-LOD confidence interval, with the peak marker indicated by a horizontal line. *A priori* hypothesized loci are marked as ellipses.

Because *S-RNase* has been hypothesized, and shown, to be a significant contributor to the UI response in some previous studies in *Solanum* (Chalivendra, *et al*., 2013; Covey, *et al*., 2010; Tovar-Méndez, *et al*., 2014), we assessed a subset of F_2_ individuals (n = 22) for expression of this protein in mature unpollinated styles. This subset represented individuals with the most and least rapid UI responses in our F_2_ population. We found that individuals that expressed S-RNase protein had a significantly more rapid UI response (n = 9, mean = 0.0815 of style) against *S. lycopersicum* pollen, compared to those that did not express *S-RNase* (n = 13, mean = 0.227 of style) (t(19.885) = 3.374, p = 0.003; **Figure 3**), indicating that S-RNase protein presence/absence is a major contributor to observed quantitative variation in UI. In comparison, all individuals were found to express HT-protein in their styles; therefore the presence/absence of HT-protein is not implicated in the phenotypic variation in UI segregating in our F_2_ population. (Note that *HT* expression may still be required for the expression of UI; see Discussion.) The F_1_ expressed both *HT* and *S-RNase* and had a rapid UI response.

**Figure 3.**
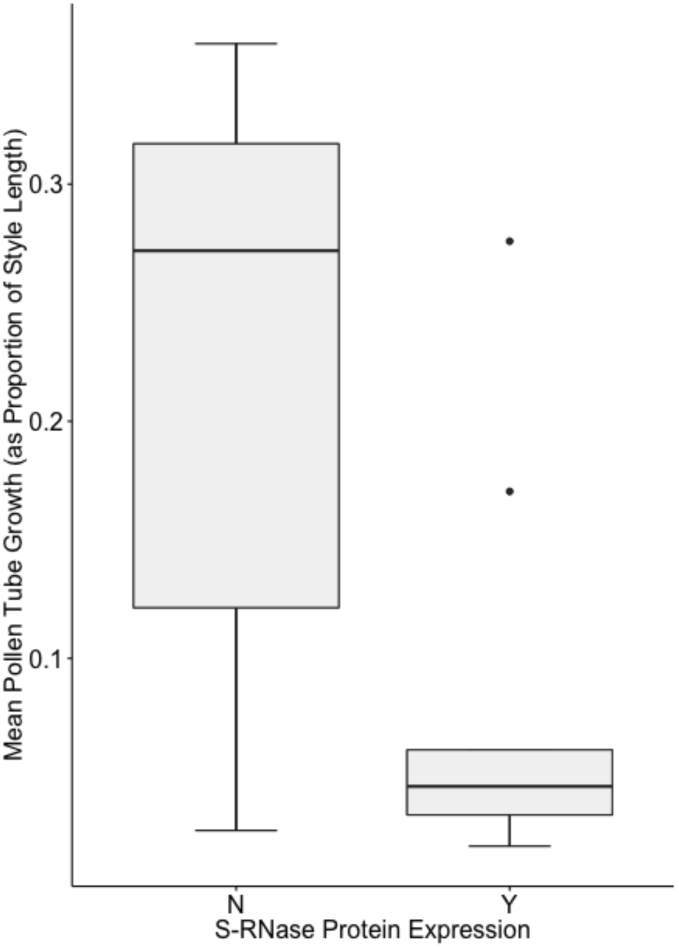
Expression of S-RNase in F_2_ mature styles as determined by a protein blot. There is significantly more rapid UI response in those styles that express S-RNase compared to those that do not (t= 3.374, p = 0.003).

Previous work in a different *Solanum* species cross also identified UI QTL on chromosomes 3 and 12 (Bernacci and Tanksley, 1997) as did a recent analysis of UI between *S. pennellii* (LA3778) and *S. lycopersicum* (LA3475) (Jewell 2016). We did not detect QTL at either of these positions. As HT-protein is thought to likely underlie the chromosome 12 UI QTL detected in other studies (Bernacchi and Tanksley, 1997; Tovar-Méndez, *et al*., 2014; Jewell 2016) this finding is consistent with our observation (above) of no differential protein expression of *HT* in our F_2_ population.

### Floral traits vary with mating system differences between the parental genotypes

The parent accessions differed in four of the six floral traits measured: stigma exsertion, style length, and ovary height and width (**Table 1**). There were no significant differences between the parent populations for corolla diameter or anther length (**Table 1, Supplementary Figure 1**). Despite large parental differences, we identified only one QTL affecting floral morphology (**Table 2**). This locus, on chromosome four, had a moderate to large effect on stigma exsertion (percent variance explained = 17.1; percent parental difference explained = 48.0). No significant QTL were detected for corolla diameter, anther length, style length, ovary height, or ovary width. Because of the limited size of the mapping population, our analyses likely missed smaller effect loci that contribute to observed parental variation in these floral traits.

### Few fertility QTL detected

We identified two QTL for fertility traits, both of which were for fruit height (**Table 2, Figure 2**) and were of small to moderate effect, explaining ∼6-16% of the variation among F2s and 5% of the parental difference each. There was no evidence for an interaction between these QTL. Interestingly, these two QTL have opposing effects on fruit height, consistent with little difference between the parental accessions in this fruit trait. Neither of the detected fertility trait QTL colocalized with our UI QTL on chromosome 1.

## Discussion

Standing genetic variation across populations within species can contribute to differences in the strength of interspecific isolating barriers. Understanding the nature of this genetic variation can provide insight into the evolutionary dynamics that shape the accumulation of these barriers among diverging species. Here we assessed the genetic basis of intraspecific variation in the strength of interspecific pistil-side unilateral incompatibility (UI). One goal was to assess whether variation in key components of mating system variation (including genetic self-incompatibility) influences this phenotypic variation. We found one large effect QTL underlying variation among populations in the rate at which UI is expressed against a second species. This QTL overlaps a major player in the self-incompatibility response--the *S-* locus--and we found that the presence/absence of stylar S-RNase protein is significantly associated with the rapidity of the UI response. Although we observe trait and genetic differences in floral and fertility traits between these two *S. pennellii* populations, some of which are typically associated with mating system transitions, QTL underlying these differences are not associated with the major effect locus controlling variation in the strength of UI. Our analysis suggests that standing variation for mate choice loci--in this case, to prevent self-fertilization--can directly affect variation in interspecific isolation--in this case a postmating prezygotic reproductive barrier.

### UI genetic mechanisms are associated with mating system loci

Both our QTL mapping analysis and our protein expression assay support the inference that S-RNase protein plays a major causal role in the observed quantitative variation in UI responses. In Solanaceae, loss of SI often involves the loss of pistil *S-RNase* expression as one of the first causal changes, so that individuals no longer reject conspecific pollen with which they share the functional pollen-side component of the SI mechanism. Our analysis indicates that this loss of pistil S-RNase protein in the SC *S. pennellii* accession (LA0716) has pleiotropic consequences for the rate at which this genotype rejects heterospecific pollen. While all individuals expressed UI within our F_2_ population, the speed of UI pollen rejection was significantly decreased when S-RNase protein was absent. Thus, we infer that standing variation within *S. pennellii* at a major mating system locus also directly contributes to how rapidly an interspecific postmating prezygotic barrier is expressed.

In addition, our data also imply that other molecular factors beyond *S-RNase* also contribute to UI expression in this species. That is, while loss of functional S-RNase protein reduced the speed of UI it did not abolish this response, suggesting other functional UI elements are retained in the pistil. S-RNase-independent UI mechanisms have been described previously in *Solanum* (Murfett, *et al*., 1996; Eberle, *et al*., 2013; Tovar-Méndez, *et al*., 2017; reviewed in Bedinger *et al*., 2017). In particular, HT protein has been implicated as a necessary molecular component of the UI response in *Nicotiana* and in other *Solanum* crosses (Bernacci and Tanksley, 1997; Covey, *et al*., 2010; Hancock, Kent and McClure, 2005; McClure, *et al*., 1999; O’Brien, 2002; Tovar-Méndez, *et al*., 2017). In addition, both SC and SI *S. pennellii* populations have previously been shown to express *HT* in their styles (Chalivendra, *et al*., 2013, Pease et al. 2016), and here we found that all tested F_2_ individuals also expressed *HT*. Together, these observations suggest that the observed quantitative variation in UI expression depends on variation in the functionality of *S-RNase*, on a background of functional *HT* expression. In addition to *HT*, there might also be other S-RNase-independent factors supporting UI function in these accessions. For example, other QTL studies have identified a major effect UI QTL on chromosome 3 in populations generated between *S. lycopersicum* and SI *S. habrochaites* (Bernacci and Tanksley, 1997) and between *S. lycopersicum* and the SI *S. pennelii* parent genotype used in our cross here (Jewell 2016, Hamlin et al. 2018). Furthermore, loss-of function of HT in SC *S. pennellii* accession (LA0716) resulted in tomato pollen rejection further down the pistil, suggesting that in addition to the S-RNAse-independent UI factors there are also HT-independent UI factors (Tovar-Méndez, *et al*., 2017). The potential contribution of additional factors to *S. pennellii* intraspecific variation in UI is testable in the future, once the specific identity of these factors is known.

Regardless, our observations support a mechanistic, explicitly genetic, association between SI and UI, consistent with other studies in closely related species (Broz et al. 2017, Tovar-Méndez, *et al*., 2014; Tovar-Méndez, *et al*., 2017). These findings in turn imply that factors governing the maintenance or loss of SI can have collateral effects on the expression of UI barriers among species. Ecologically, the loss of SI is often associated with strong selection for reproductive assurance in low density or marginal environmental conditions, where small population sizes severely restrict the availability of mating partners with different *S-*haplotypes. Under these conditions, individuals that are physiologically capable of selfing—due to loss of proteins involved in self-recognition and rejection—have a demographic and mating advantage. If SI and UI share underlying genetic mechanisms, these specific ecological conditions will also strongly determine the emergence and distribution of UI mating barriers between lineages, as will the physiological constraints governing the genetic progression of SI to SC transitions, as we discuss further below.

### No evidence for an association between UI and floral trait loci across mating transitions

Another possibility we examined was whether the strength or expression of UI was associated with other, non-SI, transitions that accompany mating system differences. Although losing SI permits selfing, the transition from facultative to predominant self-fertilization often involves additional morphological changes, especially in floral traits that affect pollinator attraction and the likelihood of self-pollination. While outcrossing species typically have larger flowers and greater distances between the receptive stigmatic surface of the female pistil and the male pollen-bearing anthers (i.e. greater stigma exsertion) (Brunet and Eckert, 1998; Motten and Stone, 2000; Rick, Holle and Thorp, 1978; Takebayashi, Wolf and Delph, 2005), highly self-pollinating species tend to have smaller flowers and smaller or no stigma exsertion (Lloyd and Barrett, 1996; Sicard and Lenhard, 2011). It is unclear to what extent changes in loci directly involved in the breakdown of SI (i.e. *S-*locus and its modifiers) work in conjunction with the genes controlling these morphological changes, as well as whether these morphological loci are associated with the expression of UI. Here, we found some floral morphology differences between the LA3778 and LA0716 parent populations that are typical of mating system differences, specifically greater stigma exsertion and greater style length in the SI compared to the SC parent genotype (Table 1), and we identified one QTL for stigma exsertion (Table 2). However, this QTL did not coincide with the UI QTL (or with known loci involved in SI). These findings provide little evidence for strong phenotypic or genetic associations between the loss of gametophytic SI and/or the expression of UI, and the morphological shifts that typically accompany mating system transitions.

### The progression to UI among species, and the role of mating system transitions

Finally, our findings also contribute to an emerging picture of the evolution of UI between species, and the specific role of mating system transitions in the formation of this post-mating prezygotic isolation barrier. First, in conjunction with mapping studies (Bernacchi and Tanksley 1997, Jewell 2016) and crossing analyses (e.g. Broz et al. 2017; see further below) in other closely-related *Solanum* species, we infer that quantitative transitions from UI competence to its loss are often associated with the cumulative loss of one or more loci functionally involved in self-incompatibility. For example, *S. habrochaites* (the sister species to *S. pennellii*) is generally an SI species within which some populations have transitioned to SC (Martin 1961), often (but not always) via the loss of functional *S-RNase* genes (Broz et al. 2016, Markova et al. 2016). Unlike the two populations of *S. pennellii* examined here, several of these *S. habrochaites* SC populations have also lost the ability to reject certain heterospecific pollen— consistent with the loss of UI competence (Baek, *et al*., 2015; Covey, *et al*., 2010, Broz et al. 2016) via the loss of one or more additional pistil-side UI factors. This greater loss of UI competence, which may be related to a longer history of self-compatibility in these *S. habrochaites* populations, appears to involve at least one other S-RNase-independent molecular player apart from *HT*, as all but one of these populations continue to express *HT* (Baek, *et al*., 2015; Covey, *et al*., 2010, Broz et al. 2017). Nonetheless, HT can clearly also contribute to this transition from UI to non-UI styles, as Tovar-Méndez, *et al*., (2017) showed that suppressing *HT* expression in *S. habrochaites* LA0407 completely abolished UI against (SC) *S. lycopersicum* pollen. Together, these observations suggest that populations undergoing a progressive loss of pistil-factors can proceed stepwise from rapid UI against other SC species (coincident with a fully functional SI system), through a transitional period of quantitative reductions in the strength of UI (coincident with the loss of pistil-side factors during the transition from SI to SC), to loss of UI against other species (coincident with the loss of both S-RNase-dependent and -independent rejection mechanisms).

Notably, while the progressive loss of pistil-side factors ultimately results in genotypes unable to mount a pistil-side UI rejection response, observations also indicate that transitions from SI to SC can be accompanied by the gain of UI, specifically the gained ability of SI lineages to reject the pollen of SC lineages (e.g. Martin 1961, Broz et al. 2016, Marcova et al. 2016). However this emergence of UI against SC lineages is not due to pistil-side changes, but instead to a progressive loss within these recently derived SC lineages of pollen-side function(s) that would otherwise neutralize pistil rejection responses (Bedinger et al. 2017). For example, within *S. habrochaites*, pollen from some SC populations is rejected by SI populations of the same species (Broz *et al*., 2017), likely due to the loss of one or more pollen-side factors specifically in these SC populations (Markova et al. 2016). In contrast, SI populations do not reject pollen from SI populations or species, indicating that SI pollen retain mechanisms for evading UI rejection in styles of other SI species. Note that the *S. pennellii* populations used here reciprocally accept each other’s pollen, indicating that the SC *S. pennellii* accession LA0716 has not progressed to the point at which UI barriers have emerged against it.

This observed progression indicates a specific temporal order to the loss of pistil-side UI and the gain of pollen-side UI rejection by other lineages, a trajectory that is strongly influenced by the dynamics governing the loss of intraspecific SI factors during transitions from SI to SC. Interestingly, in the Solanaceae (but not in other plant groups that have SI systems; Bedinger et al. 2017) this transition from SI to SC usually first involves the loss of loci that contribute to pistil-side function, and only subsequently the loss of pollen-side functions (Tovar-Méndez, *et al*., 2014). This ‘pistil-first’ transition order likely occurs because the physiology of gametophytic self-incompatibility: pollen loss–of-function mutations are incompatible on all pistils that retain pistil-side function, but genotypes with pistil-side SC mutations are able to accept all pollen donors (see also Markova et al. 2016). This leads to strong selection against pollen-side mutations because these cannot individually permit self-compatibility, and therefore will not contribute to reproductive assurance unless they are first preceded by pistil-side mutations. This expectation is supported by the observation that there are no known SC populations or species that lack SI pollen function but retain pistil function, in the Solanaceae (Tovar-Méndez, *et al*., 2014). In this way, loss of *S-RNase* and other pistil-side factors does not contribute immediately to the gain of UI, but acts as a catalyst for evolutionary changes that eventually lead to the erection of an UI barrier against the evolving population, by permitting the subsequent loss of pollen side factors. Although the conditions promoting this subsequent loss of pollen-side factors are less clear, it is possible that these mutations reduce metabolic cost (Markova et al. 2016) or increase selfing efficiency once populations have already lost pistil-side SI functions.

Regardless, it is clear that the dynamics of these mating system transitions play an influential role in the evolution of UI as a reproductive barrier. Moreover, understanding the nature of intraspecific genetic variation involved in these transitions is critical for understanding the conditions that facilitate the accumulation of reproductive isolation among populations within species (Good, *et al*., 2007; Kopp and Frank, 2005). Here, in our analysis of genetic variation within *S. pennellii*, we have shown that one of the earliest steps in this progression involves the large quantitative contribution of a pistil-side locus that is directly involved in intraspecific mate choice (via self-incompatibility). This finding agrees with previous analyses that indicate an intimate association between molecular players contributing to SI and UI. In combination with genetic and crossing data from this and other closely related species, it also suggests that intraspecific changes at these pistil-side loci are an essential antecedent step that permits the subsequent accumulation of mutations that erect new intraspecific postmating prezygotic UI barriers.

## Acknowledgements

We thank J Kissinger and DC Haak for assistance in data collection, other members of the IRBT Tomato group (Patricia Bedinger, Roger Chetelat, and Matthew Hahn) for discussions, and the Indiana University Bloomington greenhouse staff.

## Data Availability

The primary data underlying these analyses will be deposited as follows:

-morphological data: Dryad #######

-raw sequence reads and SNP genotypes: NCBI BioProject: PRJNA557135.

